# NirA is an alternative nitrite reductase from *Pseudomonas aeruginosa* with potential as an anti-virulence target

**DOI:** 10.1101/2020.12.17.423290

**Authors:** Samuel Fenn, Jean-Frédéric Dubern, Cristina Cigana, Maura De Simone, James Lazenby, Mario Juhas, Stephan Schwager, Irene Bianconi, Gerd Döring, Jonas Elmsley, Leo Eberl, Paul Williams, Alessandra Bragonzi, Miguel Cámara

**Affiliations:** National Biofilms Innovation Centre, Nottingham University Biodiscovery Institute, School of Life Sciences, University of Nottingham, Nottingham, UK; Division of Immunology, Transplantation and Infectious Diseases, IRCCS San Raffaele Scientific Institute, Milano, Italy; Present address: GLP Test Facility, San Raffaele Telethon Institute for Gene Therapy, IRCCS San Raffaele Scientific Institute, Milano, Italy; Present address: Quadram Institute Bioscience, Norwich Research Park, Norwich, Norfolk, UK; Department of Microbiology, Institute of Plant Biology, University of Zürich, Zürich, Switzerland; Present address: Institute of Medical Microbiology, University of Zürich, Zürich, Switzerland; Present address: Department of Cellular, Computational and Integrative Biology – CIBIO University of Trento, Italy; Institute of Medical Microbiology and Hygiene, University of Tübingen, Tübingen, Germany

**Keywords:** KEYWORDS, *Pseudomonas aeruginosa*, virulence target, nitrite reductase, transposon mutagenesis, genome-wide screening, disease model

## Abstract

The opportunistic pathogen *Pseudomonas aeruginosa* produces an arsenal of virulence factors causing a wide range of diseases in multiple hosts and is difficult to eradicate due to its intrinsic resistance to antibiotics. With the antibacterial pipeline drying up, anti-virulence therapy has become an attractive alternative strategy to the traditional use of antibiotics to treat *P. aeruginosa* infections. To identify *P. aeruginosa* genes required for virulence in multiple hosts, a random library of Tn5 mutants in PAO1-L was previously screened *in vitro* for those showing pleiotropic effects in the production of virulence phenotypes. Using this strategy, we have identified a Tn5 mutant with an insertion in PA4130 showing reduced levels in a number of virulence traits *in vitro*. Construction of an isogenic mutant in this gene presented similar results as those from the Tn5 mutant. Furthermore, the PA4130 isogenic mutant showed substantial attenuation in disease models of *Drosophila melanogaster*, *Caenorhabditis elegans* as well as decreased toxicity in human cell lines. This mutant also presented an 80% increased survival in murine acute and agar-bead lung infection models. PA4130 codes for a protein with homology to nitrite and sulphite reductases. Overexpression of PA4130 in the presence of the siroheme synthase CysG enabled its purification as a soluble protein. Methyl viologen oxidation assays with purified PA4130 showed that this protein is a nitrite reductase operating in a siroheme and 4Fe-4S dependant manner. The preference for nitrite and the production of ammonium revealed that PA4130 is an ammonia:ferredoxin nitrite reductase and hence was named as NirA.

## INTRODUCTION

*Pseudomonas aeruginosa* is a genetically versatile opportunistic pathogen, able to colonise and survive in multiple environments and hosts. This versatility underpins *P. aeruginosa* ability to cause a wide range of infections, commonly affecting the respiratory tract, burn wounds, urinary tract, bloodstream, cornea, skin and soft tissue (1). The majority of these infections are nosocomial, with infections in immunocompromised hosts often life-threatening.

*P. aeruginosa* has gained notoriety as a member of the ESKAPE pathogens (2). These pathogens are differentiated according to their clinical relevance and capacity to become multi-drug resistant (MDR). Often treatment of *P. aeruginosa* is unsuccessful due to high levels of intrinsic and acquired antimicrobial resistance with biofilm formation promoting antimicrobial tolerance (3, 4).

Although carbapenem-resistant *P. aeruginosa* being listed as ‘*priority one*’ by the World Health Organisation for the development of new antimicrobials, no new drugs with a novel mechanism of action against this organism have reached the market in recent years (5). Hence, there is a pressing need for the discovery of novel alternative strategies to the traditional use of antibiotics to treat *P. aeruginosa* infections. Anti-virulence therapeutic approaches have become an attractive alternative strategy to develop drugs with high specificity and narrow spectra as they reduce the illness caused by the pathogen (‘pathogen limitation’) instead of reducing pathogen burden directly (‘pathogen elimination’) (6).

In recent years, vast progress has been made on the identification of *P. aeruginosa* virulence targets, unravelling the mechanisms they employ to cause disease and developing inhibitors which can inactivate them (7–12). While these studies have uncovered numerous promising small virulence inhibitor molecules none has yet made it to the clinic. This has been influenced by many different factors, including the reliance on a single disease model, the potential conservation with the microbiota, the lack of understanding of target functionality and the inability to define success when searching for inhibitors (13).

The sequencing of the first *P. aeruginosa* genome in 2000 revealed that the PAO1 strain sequenced (PAO1-UW) has a genome size of 6.3 Mbp and contains 5570 open reading frames, making it the largest genome sequenced at the time (14). This large genome underpins the extensive metabolic and regulatory network providing *P. aeruginosa* with the genetic versatility to colonise multiple environments, hosts and host sites. Besides, with the function of only 22.7% of *P. aeruginosa* genes experimentally demonstrated and close to 2000 genes without functional annotation (15), there is still a large amount of information missing with regard to the mechanisms by which this organism causes disease, and potentially a vast array of novel *P. aeruginosa* virulence targets remains to be discovered.

Early studies suggested that virulence mechanisms employed by *P. aeruginosa* to infect phylogenetically diverse hosts are remarkably well conserved. Comparison of infection mechanisms in the plant *Arabidopsis thaliana* and mice revealed that *P. aeruginosa* uses a shared subset of virulence genes to provoke disease (16). The conserved nature of *P. aeruginosa* virulence suggested that use of a single disease model is sufficient to dissect *P. aeruginosa* virulence in all hosts (17), with various studies utilising the nematode *Caenorhabditis elegans* (18), fruit-fly *Drosphila melanogaster* (19), silkworm *Bombyx mori* (20), larvae *Galleria melonella* (21) and Zebrafish embryos (22). However, limitations of these studies are related to the impact of virulence in multi-host system and the number of *P. aeruginosa* strains tested.

A study performed by our group combining whole-genome transposon mutagenesis with a cascade of *in vitro* and *in vivo* infection models uncovered the host-specific nature of *P. aeruginosa* virulence (23). This revealed a remarkably low overlap in virulence factor requirement between models suggesting that many of the virulence factors identified with single model studies may not represent virulence factors required during human disease (23). Given the broad range of clinical manifestations exhibited by *P. aeruginosa* infections, it stands to reason that virulence determinants identified as attenuated in multiple models are more likely to be both relevant in human disease, and at multiple infection sites, representing promising anti-virulence targets.

This study builds upon the successful whole-genome transposon mutant screening to identify novel *P. aeruginosa* virulence target genes (23). Here, we describe the identification of an additional mutant from this library on a yet uncharacterised gene of *P. aeruginosa*, PA4130, exhibiting attenuation in all infection models tested. Functional characterisation revealed that PA4130 encodes an assimilatory nitrite reductase with a potential role in nitrogen source metabolism during *P. aeruginosa* pathogenesis. The attenuation in all disease models tested and the lack of human homologues make this nitrite reductase a promising anti-virulence therapeutic target.

## RESULTS

### Isolation of a *P. aeruginosa* mutant attenuated in multiple virulence factor production

To isolate novel virulence genes, a transposon (Tn5) insertion library was generated in a wild-type PAO1-L strain (23). A total of 57,360 individual colonies were picked and screened for pleiotropic attenuation in virulence phenotypes (reduced swarming, exoprotease and pyocyanin production) (23). Using the same screening approach, in the current study a further transposon mutant (PAJD21) was isolated from the library. This mutant displayed reduced levels of pyocyanin and pyoverdine production as well as decreased swarming motility, with twitching, swimming, protease and elastase activity being unaffected when compared to the wild type (**Fig. 1A, 1B, 1C, 1D** and **Table S1**). Nucleotide sequence analysis of the Tn5 flanking region showed that the transposon had inserted into PA4130, encoding a hypothetical protein with homology to nitrite and sulphite reductases, and forming a predicted operon with PA4129 (**Fig. 1E**). To ensure the attenuation in virulence traits observed is not due a polar effect on PA4129, located in the same predicted transcriptional unit as PA4130, in-frame deletion mutants of both PA4130 and PA4129 were constructed to generate strains PAJD25 and PASF06 respectively. These isogenic mutants showed no growth difference in either LB or artificial sputum media (ASM) in relation to the parental strain (**Fig. S2**).

**FIG 1.**
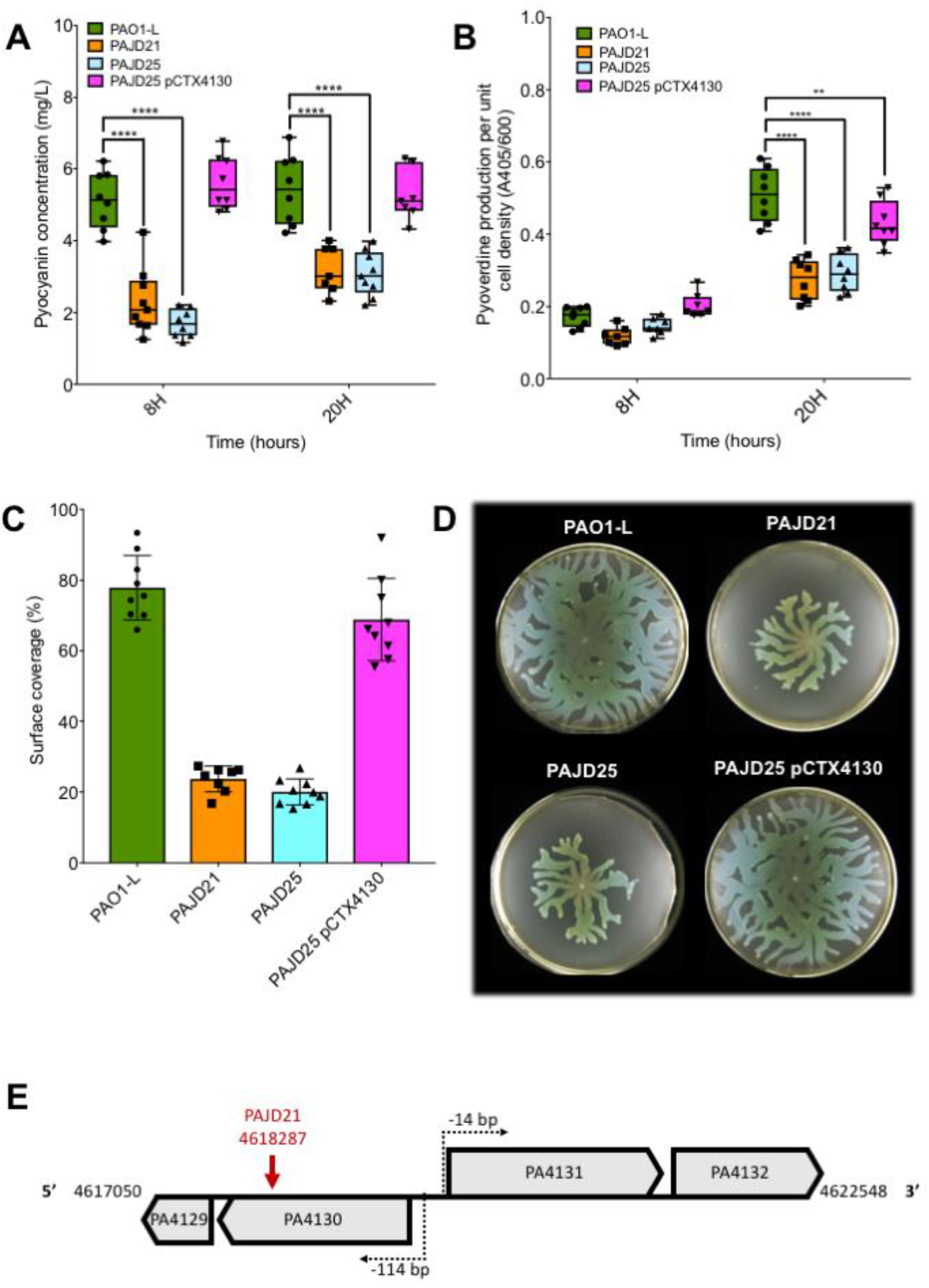
*In vitro* phenotypic characterisation of PAO1-L virulence factor production versus PA4130 mutants. The transposon mutant PAJD21 (PA4130::*Tn5*), the PA4130 in frame deletion mutant PAJD25 and the complemented mutant PAJD25 pCTX4130 were used in these experiments (A) Pyocyanin levels determined in supernatants from cultures grown for 8 and 20 hours in LB using a colorimetric assay. (B) Pyoverdine levels determined in supernatants from cultures grown for 8 and 24 hours in modified artificial sputum media using a colorimetric assay. Swarming motility surface coverage measurements (C) and images (D); cells were inoculated from a 16h LB culture onto swarming plates containing 0.5% agar and incubated at 37°C for 16 hours. (E) Diagram displaying operon structure and transcriptional start sites of PA4129-30 and PA4131-32 with the PAJD21 *Tn5* insertion site indicated by the red arrow. Data were collated from 3 independent experiments with at least 3 replicates each. **P <0.01, ****P <0.0001, Dunnett’s multiple comparisons test.

PAJD25 showed similar phenotypes to the Tn5 mutant PAJD21, with pyocyanin and pyoverdine reduced in both LB and ASM and swarming motility also impeded. Chromosomal complementation of PAJD25 with PA4130 restored pyocyanin and swarming motility to wild-type levels, with partial restoration of pyoverdine production (**Fig. 1A, 1B, 1C, 1D**). In contrast, the PA4129 deletion mutant PASF06 did not show any significant alterations in the virulence-related phenotypes (**Table S3).** This demonstrates that the phenotypes observed are specific to the mutation of PA4130.

When identifiying new virulence targets it is paramount to ensure they are conserved in a wide range of strains. To establish this, a Nucleotide BLAST of PA4130 against the NCBI *P. aeruginosa* taxonomic identifier database (taxid:287) revealed that this hypothetical protein is highly conserved, with 363/367 of the available genomes encoding a PA4130 orthologue. To determine whether PA4130 has a similar role in the virulence of *P. aeruginosa* strains from different sublines, in-frame PA4130 deletion mutants were created in the clinical isolates PA7 Bo599, PA14 AUS471 and LESB58 PA-W39. Deletion of PA4130 across all these strains resulted in a similar attenuation for both pyocyanin production and swarming motility whilst protease production and growth remained unaffected, validating the results obtained for PAJD21 and PAJD25 (**Fig. 2A,2B, 2C and Table S4**). In contrast, reduction in pyoverdine production was not conserved with no consistent pattern emerging between strains (**Table S4**). The similar phenotypes observed in the PA4130 deletion strains across phylogenetically diverse *P. aeruginosa* strains confirms PA4130 is not a PAO1-L specific virulence factor. This supported the use of PAO1-L and the derived PAJD25 as a representative strains to further characterise the role of PA4130 in virulence.

**FIG 2.**
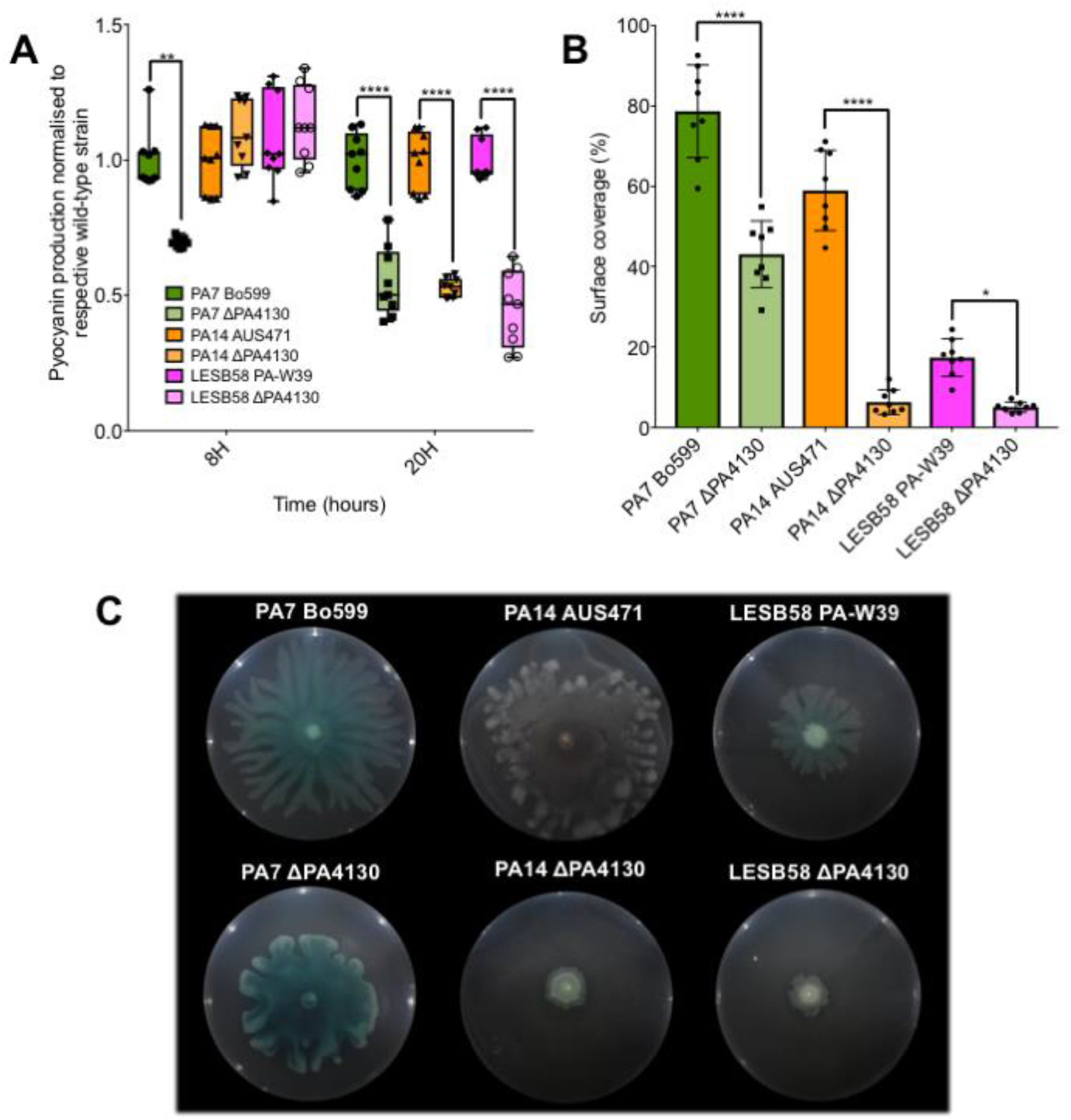
*In vitro* phenotypic characterisation of virulence factor production for PA4130 orthologue mutants in PA7 Bo599, PA14 AUS471 and LESB58 PA-W39. (A) Pyocyanin levels determined in supernatants from cultures grown for 8 and 20 hours in LB using a colorimetric assay. Swarming motility surface coverage measurements (B) and images (C); cells were inoculated from a 16h LB culture onto swarming plates containing 0.5% agar and incubated at 37°C for 20 hours. Data collated from 2 independent experiments with at least 4 replicates.*P <0.05, **P <0.01, ****P <0.0001, Tukey’s multiple comparisons test.

### PA4130 is required for full virulence in *C. elegans* and *D. melanogaster*

To establish whether the decrease in virulence trait production observed in the PA4130 deletion mutant PAJD25 *in vitro* also results in disease attenuation *in vivo*, the *D. melanogaster* and *C. elegans* non-mammalian infection models previously used for *P. aeruginosa* infection studies were initially used. In both disease models the PAJD25 (ΔPA4130) mutant showed an attenuated survival rate when compared to the isogenic wild-type PAO1-L (**Fig. 3**). For *C. elegans*, PAJD25 virulence was severely attenuated with a 57% increase in survival at 72h post-infection (**Fig. 3A**). In the case of *D. melanogaster,* PAJD25 killing was delayed with a 40% increase in survival at 18h post infection, although by 22 hours this difference was negligible (**Fig. 3A**).

**FIG 3.**
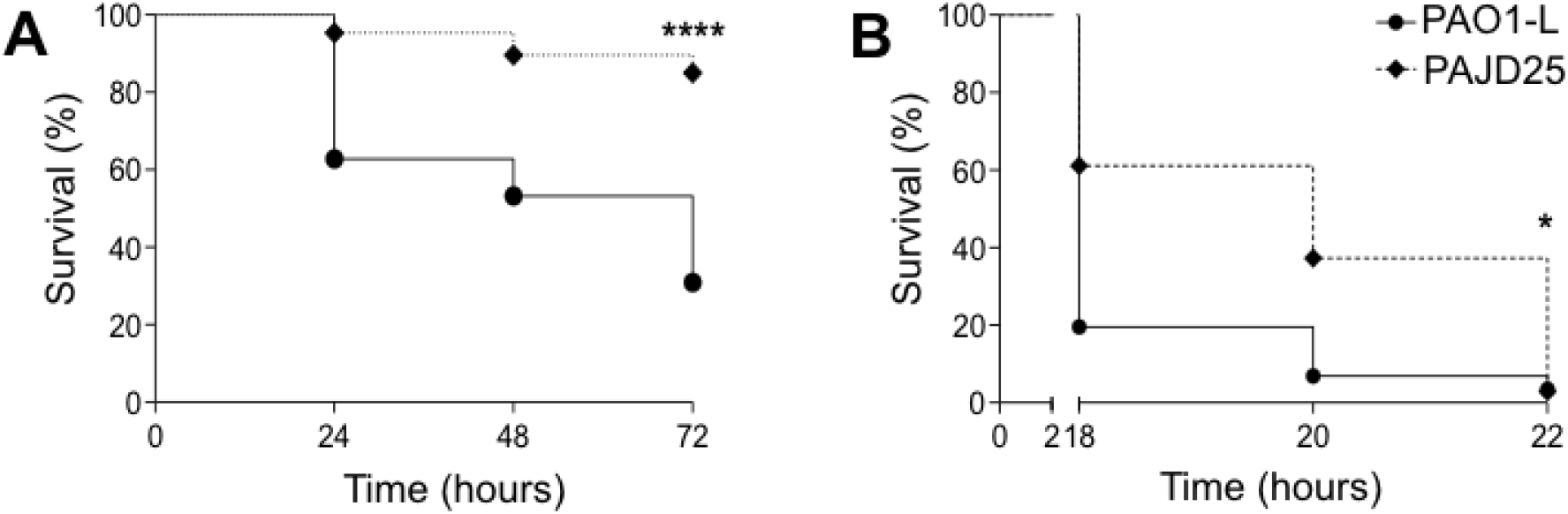
Survival of PAO1-L versus PAJD25 in non-mammalian models. (A) 72h lethality curve in the *C. elegans* infection model. (B) 22h lethality curve in *D. melanogaster*. Both models demonstrated clear attenuation with increased survival of PAJD25 in *C. elegans* at all time-points. *D. melanogaster* exhibited a delay in killing with increased survival at earlier time-points; however, by 22 hours both PAO1-L and PAJD25 showed almost complete killing. The results presented display the mean values from three independent experiments. *P <0.032, ****P <0.0001 Log rank Mantel-Cox test.

### Deletion of PA4130 results in reduced cytotoxicity, cellular invasion and IL-8 production in A549 human lung epithelial cell line

Previous work performed in the Tn5 mutant library screening by Dubern and colleagues (23) suggested that attenuation using *in vitro* assays and invertebrate infection models does not translate into mammalian models. To establish whether this is the case for the PA4130 mutant, the PAJD25 (ΔPA4130) strain was tested on the A549 pulmonary cell line for cytotoxicity, invasion and IL-8 production (**Fig. 4**). Cytotoxicity of the PAJD25 supernatant was drastically reduced compared to PAO1-L, which is in line with the reduced levels of secreted pyocyanin (**Fig. 1A**and **4A**). Furthermore, invasion of A549 epithelial cells was decreased in PAJD25, with IL-8 production also reduced when compared to wild-type strain PAO1-L (**Fig. 4B** and **4C**).

**FIG 4.**
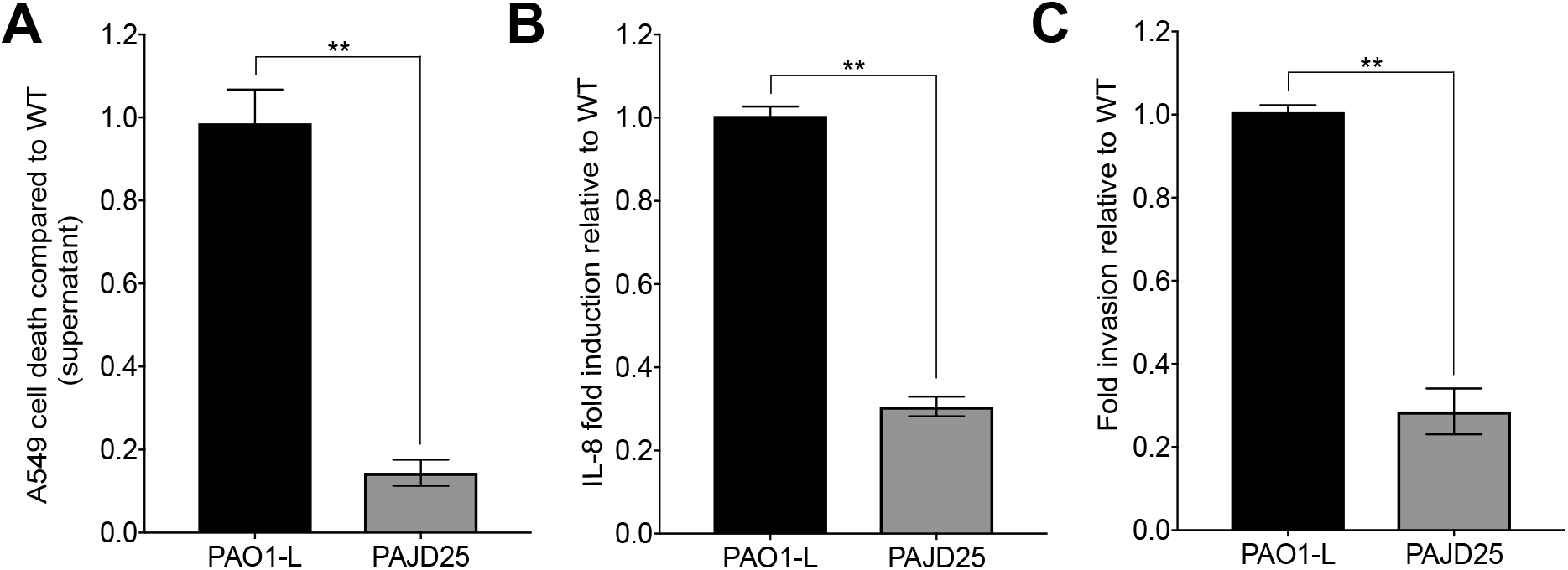
Cytotoxicity, IL-8 release and invasion of A549 cells following infection with either PAO1-L or PAJD25. (A) Cytotoxicity assayed with Syto13/PI viability staining. (B) IL-8 released quantified by ELISA following infection. (C) Invasion quantified using antibiotic (polymyxin B) exclusion assay. Data, from three independent experiments, expressed as mean (+/−) standard error of mean (SEM). **P<0.01, Student’s t-test.

### Deletion of PA4130 reduces lethality of *P. aeruginosa* in acute and agar-bead murine infection models

The impact of a PA4130 mutation on *P. aeruginosa* virulence was initially assessed by determining the survival curve of PJD25 (ΔPA4130) in an acute lung infection model in C57BL/6NCrlBR mice using an infection dose of 5×10^6^ colony forming units (CFU). A survival rate of 80% was shown at 72h post-infection in contrast to PAO1-L which did not show any survival after 36h (**Fig. 5**).

**FIG 5.**
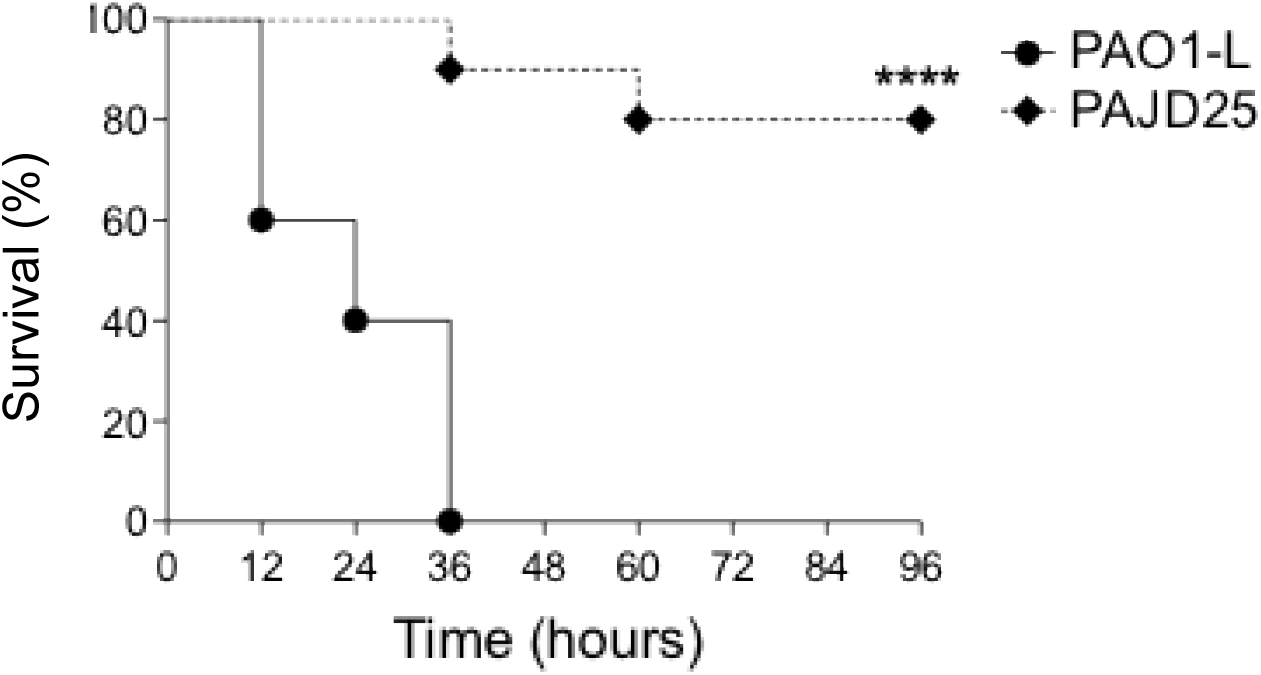
Survival curve of C57BL/6NCrlBR mice infected with PAO1-L and PAJD25 in an acute lung infection model. At 36 hours PAJD25 exhibits a 90% increase in survival when compared to PAO1-L, with 80% of PAJD25 infected mice surviving past 96 hours. ****P<0.0001, Log rank Mantel-Cox test.

The effect of the PA4130 mutation was then tested in an agar bead infection model, to monitor initial colonization and systemic spread. C57BL/6NCrlBR mice were infected with a 2×10^6^ CFU dose of either PAO1-L or PAJD25. At 18 hours post infection, 20% of PAO1-L and 80% of PAJD25 infected mice survive. By 36 hours 80% of PAJD25 challenged mice still survive whilst PAO1-L infected mice exhibit no survival (**Fig. 6A**). The reduction in mortality was confirmed by the decreased CFU recovery from the lung, liver and spleen of the mutant compared to wild type (**Fig. 6B, 6C** and **6D**). Clearance or reduced CFUs in the lung of PAJD25 infected mice demonstrates that interruption of PA4130 interferes with colonisation. This impaired colonisation results in reduced systemic spread with no PAJD25 detected in the spleen or liver in all but one sample (**Fig. 6B, 6C** and **6D**).

**FIG 6.**
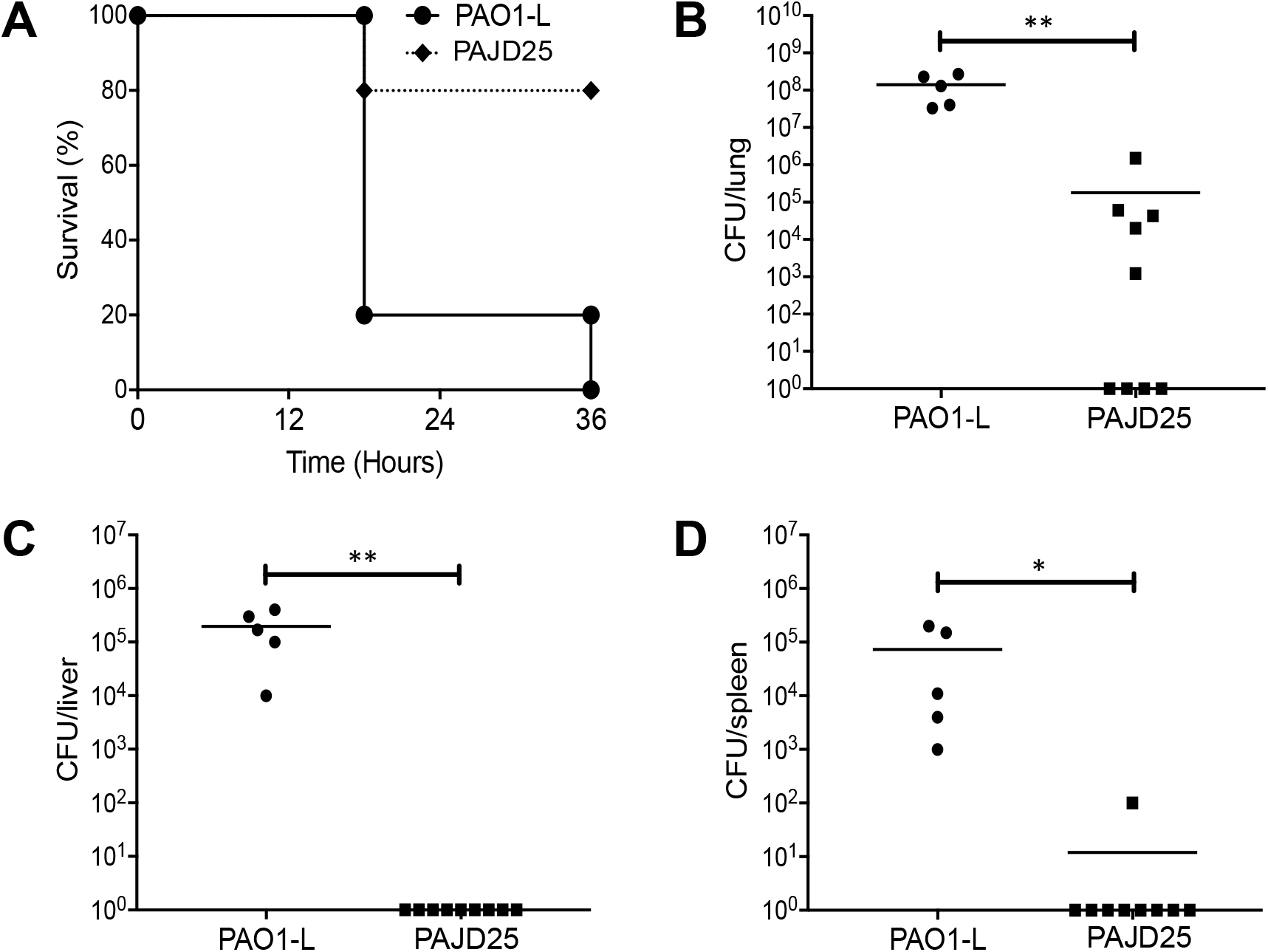
Virulence of PAO1-L and PAJD25 in an agar bead murine lung infection model of colonisation in C57BL/6NCrlBR mice. (A) Survival curve, (B) CFU recovery from lung, (C) CFU recovery from the liver and (D) CFU recovery from the spleen at 36 hours post infection. PAJD25 exhibits a significant increase in survival at 36h post inoculation with a CFU recovery 3-log lower than PAO1-L in the lungs. PAO1-L infection progresses to the liver and spleen within 36h whilst PAJD25 infection does not proceed to the liver and spleen as indicated by the absence of CFU recovery in all but one sample. ****P <0.0001 Log rank Mantel-Cox test for survival curve. *P <0.05, **P < 0.01 Student’s t-test for CFU recovery.

Overall, the data presented demonstrates that PA4130 plays a role in *P. aeruginosa* survival and virulence in multiple models of infection, suggesting that PA4130 is not a model-specific virulence gene.

### Purification of PA4130 reveals characteristics of a possible nitrite or sulphite reductase

Amino acid sequence analysis showed that PA4130 has an amino acid similarity of 21% with the *E. coli* CysI sulphite reductase hemoprotein subunit and that no protein homologues are encoded within the human genome. Alignment of the PA4130 amino acid sequence with the *E. coli* MG1655 CysI, *Mycobacterium tuberculosis* H37RV NirA and *Spinecea oleracea (spinach)* NirA, using ClustalW, revealed the presence of conserved residues indicative of 4Fe-4S and siroheme prosthetic group binding sites (**Fig. S2**). Initial expression and purification attempts yielded insoluble, non-functional PA4130. Prosthetic group limitation was hypothesized to limit soluble PA4130 expression. Co-overexpression of iron-sulphur cluster biogenesis or siroheme synthesis has been demonstrated to increase expression of functional enzymes incorporating these prosthetic group (24, 25).

PA4130 was subsequently co-overexpressed in the presence of the siroheme synthase CysG, yielding a 20-fold increase in soluble PA4130 expression. PA4130 was purified to homogeneity with a combination of immobilised metal affinity chromatography and chitin column chromatography, whilst size exclusion chromatography revealed PA4130 is a monomer in solution (**Fig. S3).** Spectroscopic characterisation of the soluble PA4130 protein confirmed the presence of peaks at 382nm, 587nm and 712nm, characteristic of siroheme and 4Fe-4S incorporation into nitrite/sulphite reductase hemoprotein sub-units (data not shown).

### PA4130 codes for an assimilatory nitrite reductase operating in a ferredoxin dependant manner

Nitrite or sulphite reductases catalyse the 6-electron reduction of nitrite or sulphite to ammonium or hydrogen sulphide, respectively. These enzymes require an electron acceptor and donor to complete the electron transport chain. The electron acceptor is nitrite or sulphite with the electron donor being either NADPH, NADH or ferredoxin.

To determine the native electron acceptor, we used a reduced methyl viologen (MV) reductase assay, which spectrophotometrically tracks the artificial electron donors oxidation in the presence of nitrite or sulphite as the possible electron acceptors for PA4130 (26). Our data show that PA4130 was only able to oxidise MV in the presence of nitrite with minimal oxidation occurring in the presence of sulphite (**Fig. 7A**). These results confirm that PA4130 functions as a nitrite reductase and hence participates in the nitrate reduction pathway.

**FIG 7.**
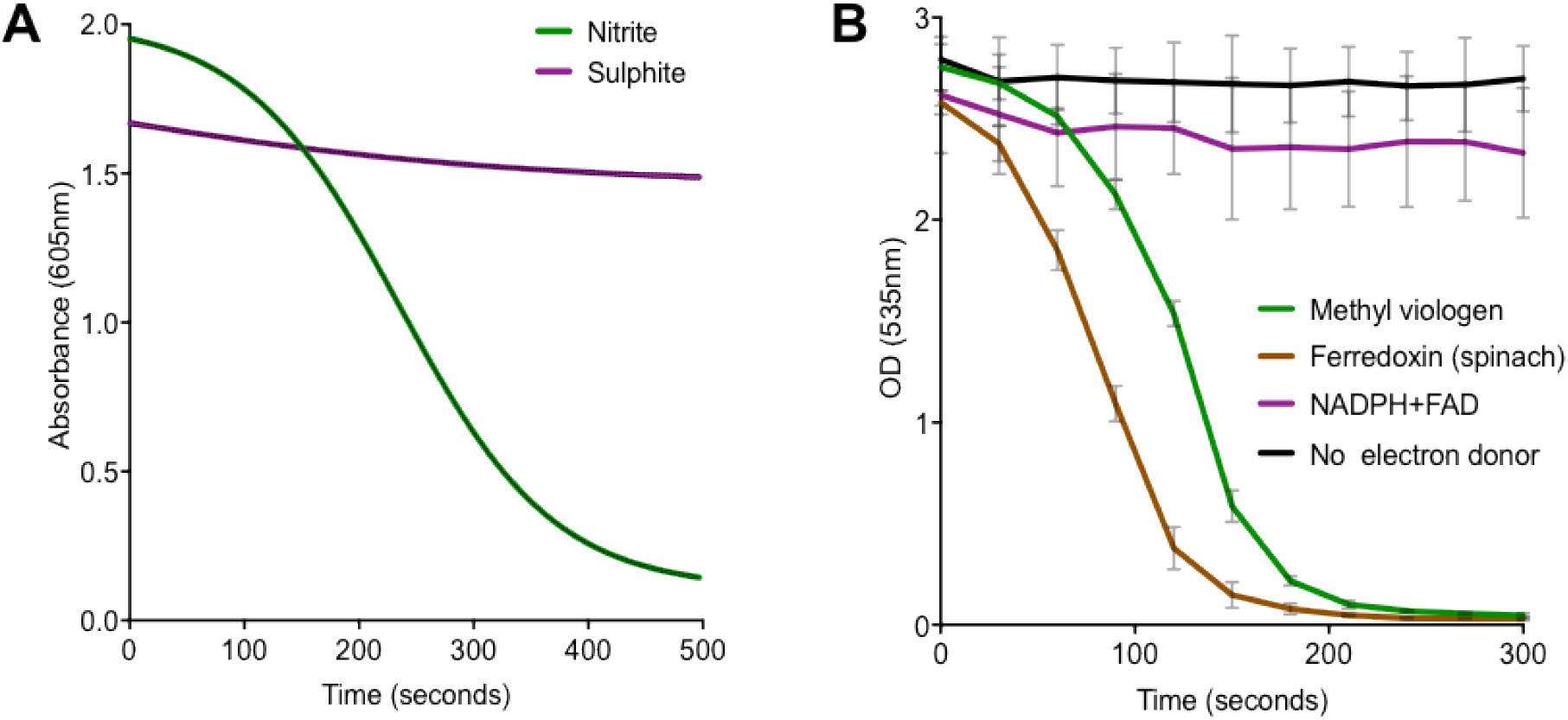
PA4130-mediated reduction assays for native electron acceptor and donor determination. (A) Methyl viologen oxidation assay in the presence of the electron acceptors nitrite and sulphite. Reduced MV is only oxidised in the presence of nitrite indicating PA4130 is a nitrite reductase. (B) Nitrite reduction assay using MV, reduced ferredoxin and NADPH as electron donors. Remaining nitrite quantified by Griess diazotization, with nitrite reduction occurring in the presence of MV and reduced ferredoxin.

To identify the native electron donor used by PA4130, the same assay as described above was carried out with the alternative electron donors NADPH+FAD and reduced spinach ferredoxin, with remaining nitrite quantified using Griess diazotization. Tracking of the remaining nitrite revealed that reduction only occurs in the presence of MV and reduced ferredoxin whilst minimal reduction occurs with NADPH+FAD (**Fig. 7B**). This observation is in agreement with a study from Frangipani and colleagues showing that increased expression of both PA4130 and the ferredoxin:NADP+ reductase *fprA* are induced by cyanogenesis in *P. aeruginosa* (27). This suggests that the role of FprA is to reduce ferredoxins, using NADPH as an electron donor, ensuring a supply of reduced ferredoxin for PA4130 to derive electrons for nitrite reduction

The nitrite reductase cognate pathways can be assigned via the end reaction product. Nitric oxide (NO) forming nitrite reductases participate in the dissimilatory denitrification pathway whilst ammonium (NH4+) producing reductases are part of either assimilatory or dissimilatory nitrite reduction to ammonium (DNRA) pathways (**Fig. 8**) (28). *P. aeruginosa* encodes 3 structurally distinct nitrite reductases, the NO-forming denitrification reductase NirS, NH_4_ ^+^ forming NirBD and the unassigned PA4130. Conserved domain analysis using DELTA-BLAST suggests that PA4130 resembles an ammonium producing nitrite reductase. This was confirmed with an ammonium production assay following nitrite reduction using reduced ferredoxin as an artificial electron donor (**Fig S4**).

**FIG 8.**
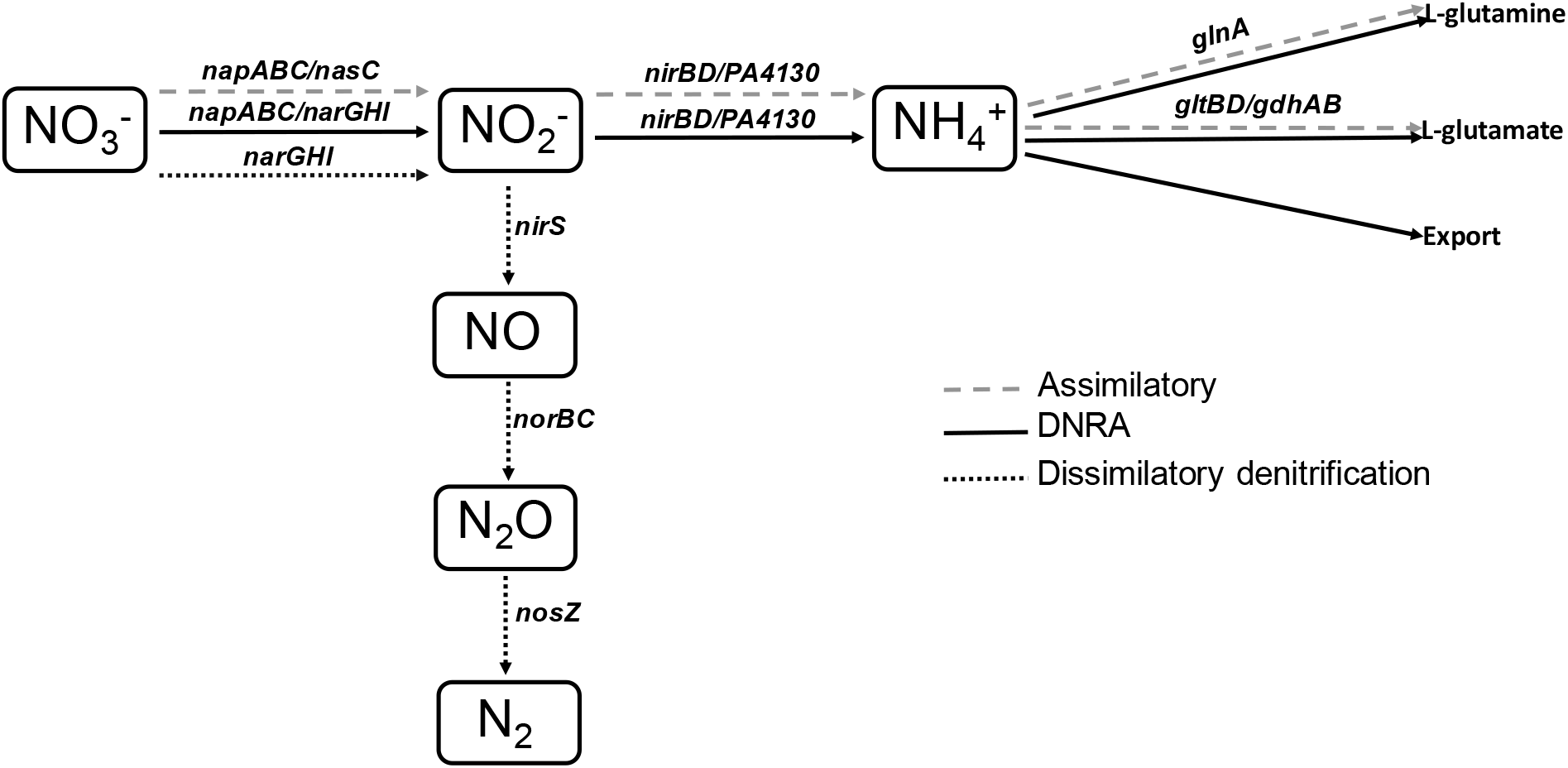
Genetic and biochemical pathways involved in nitrate reduction. This highly branched process is divided into three primary pathways: assimilatory, dissimilatory nitrite reduction to ammonium (DNRA) and dissimilatory denitrification. The presence of multiple pathways enables multiple biological functions to be performed with amino acid synthesis, detoxification, energy generation and conservation key to survival in diverse environments.

Together these data indicate that PA4130 codes for an ammonium producing, ferredoxin and siroheme-dependant, nitrite reductase, which participates in the nitrate reduction cycle. We propose that PA4130 be renamed NirA to fit with current nomenclature.

## DISCUSSION

In a preceding study, we aimed to identify novel genes required for virulence in *P. aeruginosa* using a multi-host screening strategy with the ultimate goal of discovering novel anti-virulence targets (23). Using the same strategy we have identified a new Tn*5* mutant in PA4130 which is attenuated in multiple virulence factor production. One limitation previously identified with this screening strategy (23) was the use of a single strain of *P. aeruginosa* as there can be significant variations between the virulence of different strains, potentially leading to the identification of strain-specific virulence factors (29). This problem was circumvented by generating PA4130 orthologue deletion mutants in phylogenetically distinct *P. aeruginosa* strains covering the major phylogenetic groups identified by Freschi and colleagues (30). This revealed a conserved role of PA4130 in virulence factor production with pyocyanin production and swarming motility being impaired across all mutants.

Further screening revealed attenuation of a PA4130 mutant across all disease models tested in this study, confirming that PA4130 is not a disease model specific virulence factor and is likely to play a key role in survival during human infection. This makes PA4130 a promising target which can be taken forward for functional and structural characterisation with a view to develop novel virulence inhibitors against this opportunistic pathogen.

Characterisation of the predicted gene product of PA4130 showed an assimilatory ferredoxin dependant nitrite reductase likely involved in nitrate and the wider central nitrogen metabolism. Hence, it has subsequently been named as *nirA*. Previous studies have also detected various carbon and nitrogen metabolic genes as essential for full virulence since infection is associated with significant metabolic changes (23, 31, 32). Inhibiting the ability to adapt to carbon and nitrogen sources availability during infection could prove an attractive anti-virulence strategy with metabolic dysregulation having knockdown effects on multiple cellular cycles.

Nitrogen source metabolism is essential for the survival of *P. aeruginosa* in diverse environments enabling amino acid synthesis, carbon source utilisation and respiration in the absence of oxygen (31). A key element to nitrogen source metabolism is the reduction of nitrate. *P. aeruginosa* nitrate metabolism is a highly branched and interlinked process with three main pathways, dissimilatory denitrification, dissimilatory ammonification and assimilatory (**Fig. 8**) (28, 33).

Denitrification reduces nitrate to gaseous dinitrogen. The enzymes involved in this reaction form a redox-loop with respiratory dehydrogenases allowing the generation of a proton gradient, when coupled with electron transfer into the quinone pool via the dehydrogenase (28, 33). This facilitates respiration in the absence of oxygen. Assimilatory and dissimilatory nitrate reduction are similar in that the final product produced is ammonium utilising the NADH-dependant assimilatory nitrite reductase NirBD (34, 35). The assimilatory pathway is dedicated to biosynthesis, with the assimilatory nitrate reductase (NasC) and assimilatory nitrite reductase (NirBD) producing nitrogen in the form of ammonium. This can be used as a nitrogen source for growth, primarily via production of L-glutamine and L-glutamate. Dissimilatory ammonification reduction differs, requiring the respiratory nitrate reductases NarGHI or NapABC to enable energy generation/conservation (28, 33) whilst simultaneously maintaining intracellular level of nitrogen via the production of ammonium.

The NirA production of ammonium *in vitro* suggests that it participates in either assimilatory or dissimilatory nitrate reduction with potential implications in amino acid biosynthesis, energy conservation, detoxification and maintenance of the intracellular redox environment (28, 34). However, determining the exact role NirA plays during virulence is complicated by the fact that *P. aeruginosa* codes for an additional ammonium producing nitrite reductase NirBD.

Although NirBD and NirA perform the same molecular function, they are differentially regulated. *NirBD-PA1779(nasC)-cobA* are under control of the RpoN nitrogen utilisation sigma factor with NtrC and NasT acting as transcriptional activators. Expression occurs under low nitrogen availability in the presence of nitrate and nitrite (35). In contrast, NirA expression is under control of cyanogenesis and is involved in protecting cells from HCN self-intoxication alongside PA4129-34 and *cioAB* by a yet undefined mechanism (27). The role played by these ammonium-forming nitrite reductases during pathogenesis has largely remained unexplored *in vivo* with the exception of *M. tuberculosis,* where induction of *nirBD* is required for survival in human macrophages during hypoxia (36). Differential regulation of NirA is likely to be key to the virulence attenuation seen in strain PAJD25 in the different disease models tested in this study. Production of HCN has been demonstrated to be essential for *P. aeruginosa* virulence in multiple models (37–39) and is subject to control by the anaerobic regulator ANR and the *las*/*rhl* quorum sensing circuits (40). This ensures that HCN maximally occurs under microaerophilic conditions, limiting inhibition of aerobic and anaerobic respiratory chains (40). With *nirA* expression shadowing HCN production we hypothesize that NirA may play a similar role to NirBD of *M. tuberculosis*, in protecting *P. aeruginosa* under low oxygen conditions during cyanogenesis. Whether the NirA specific mechanism is protective, through the detoxification of nitrite, or biosynthetic in the form of ammonium production, remains to be determined. Further work is required to further characterise ammonium-forming nitrite reductases and the overlapping*, in vivo* roles they may play during *P. aeruginosa* pathogenesis.

Whilst the mechanisms are yet to be unravelled, the discovery that the nitrite reductase NirA is required for virulence in multiple infection models represents a very attractive prospect for the development of inhibitors. Humans do not reduce nitrate and nitrite and as such do not encode nitrite reductases, minimising off-site effects of any inhibitors developed. Taking these points together makes NirA a promising anti-virulence target candidate against *P. aeruginosa* infections.

## MATERIALS AND METHODS

### Bacterial strains, plasmids, growth conditions and DNA manipulation

Bacterial strains and plasmids are listed in table S1. *P. aeruginosa* and *E. coli* were routinely cultured in Lysogeny broth (LB) at 37oC with vigorous shaking (200rpm), unless otherwise stated. Artificial sputum media (ASM) was made according to (41), modified with the addition of 0.5mM KNO_3_ to replicate conditions reported in the CF lung (42, 43). Media were solidified with 1.5% agar. Antibiotics were added, when required, at concentrations of 20 μg ml^−1^ for gentamicin, 15 μg ml−1 for nalidixic acid and 20 μg ml^−1^ for streptomycin. Tetracycline was added at a final concentration of 150 μg ml−1 for *P. aeruginosa* and 10 μg ml^−1^ for *E. coli*. Protein overexpression was performed in Terrific broth (TB) (24 g/L yeast extract, 12 g/L tryptone, 4% glycerol, 0.017M KH_2_PO_4_, and 0.072 KH_2_PO_4_) supplemented with 1mM FeSO_4_.6H_2_0 and 50 μg ml^−1^ of carbenicillin and streptomycin.

Genomic DNA isolation was performed using a Wizard Genomic DNA purification kit (Promega). Plasmid isolation was performed with GenElute plasmid miniprep kit (Sigma). All other standard DNA manipulation techniques such as analysis, digestion, ligation and transformation were performed according to Sambrook and Russell (44).

### Mutant selection and phenotype confirmation

A previously reported *P. aeruginosa* Tn5 mutant library was further screened for new mutants showing alterations in pyocyanin production, swarming motility and alkaline protease activity with the aid of Flexis (Genomics solutions) colony picking robot as previously described (23). Quantitative assays confirming observed phenotypes for pyocyanin, pyoverdine, swarming and protease were evaluated as described elsewhere (45).

### *C. elegans* and *D. melanogaster* virulence assays

Both Nematode slow killing assays and *D. melanogaster* disease models were performed as previously described (23, 46).

### Cell culture, IL-8 secretion, invasion and cytotoxicity assays

A549 (human type II pneumocytes) were purchased from ATCC and cultured as described (47). IL-8 secretion was performed as reported previously (23). Bacterial invasion was determined using a polymyxin B protection assay (23). Cytotoxicity of *P. aeruginosa* culture supernatant and cell pellet was assessed using the Syto-13/propidium iodide viability test according to (48).

### In-frame deletion mutant construction and complementation

To construct an in-frame deletion in PA4130, two DNA fragments 427bp upstream and 433bp downstream from PA4130 were generated and fused by overlap extension PCR using PAO1-L genomic DNA as a template. The upstream 427bp fragment was amplified with primers PA4130F1 which carries an XbaI restriction site and PA4130R1 containing the first 12 nucleotides of PA4130 with an overhanging end containing the last 15 nucleotides of the PA4130 ORF (**Table S1**); the downstream 433bp fragment was amplified with PA4130F2 containing the last 15 nucleotides of PA4130 with an overhanging end containing the first 12 nucleotides and PA4130R2 containing a HindIII restriction site (**Table S1**). To perform the overlap extension PCR a secondary PCR was performed with the 427bp and 433bp fragments serving as templates and primers PA4130F1/PA4130R2 (**Table S2**). The final PCR product was cloned using the XbaI/HindIII restriction sites into the vector pME3087, resulting in the suicide plasmid pME4130. The suicide plasmid used to generate the PA4129 mutant was construct as above, using primer pairs PA4129F1/PA4129R1 and PA4129F2/PA4129R2 to generate two PCR products upstream and downstream of PA4129 and using primer pairs PA4129F1/PA4129R2 to generate the final PCR product containing a deletion in PA4129, which was cloned into pME3087 resulting in the suicide plasmid pME4129 (**Table S2**).

The PA4129 and PA4130 in frame deletions were generated by allelic exchange using pME4129 and pME4130 respectively. Briefly, pME4129 and pME4130 was mobilised by conjugation into the relevant *P. aeruginosa* strains using *E. coli* S17.1 *λpir*. Conjugants were selected for on LB agar supplemented tetracycline and nalidixic acid. Strains were re-streaked twice on LB, with no antibiotic, and subjected to tetracycline sensitivity enrichment to select for double cross-over events (48). Colonies were screened for loss of resistance to tetracycline with allelic exchange confirmed with PCR and DNA sequencing, resulting in strains PASF06 (ΔPA4129) and PAJD25 (PAO1-L ΔPA4130), PA7 Bo599 ΔPA4130, PA14 AUS471 ΔPA4130 and LESB58 ΔPA4130 (**Table S2**).

PAJD25 was complemented by amplifying a 2172bp fragment containing the PA4130 ORF and +498bp of the translational start site using primers 4130CTXF1/4130CTXR1. The PCR product was cloned into the integrative vector mini-CTX-1 using the HindIII/BamHI restriction sites forming pCTX4130, with the resulting vector mobilised by conjugation, as performed with pME4129 and pME4130. Conjugants were selected on LB agar supplemented with tetracycline and nalidixic acid. The integration of pCTX4130 was confirmed using PCR and DNA sequencing with complementation demonstrated through restoration of pyocyanin production and swarming motility.

### Acute murine infection model

C57BL/6NCrlBR male mice (8-10 weeks of age) were purchased from Charles River Laboratories, Italy. In the acute murine lung infection model, *P. aeruginosa* strains were grown for 3h in tryptic soy broth (TSB). Bacteria were then harvested, washed twice with sterile PBS and the OD of the bacterial suspension adjusted to an OD_600_ nm. Planktonic bacteria were re-suspended in sterile PBS to the desired dose for infection of 5×10^6^ CFUs/mouse. Mice were anaesthetized and the trachea directly visualised by a ventral midline incision, exposed and intubated with a sterile, flexible 22-gr cannula attached to a 1 ml syringe accordingly to established procedures (47). A 60ul inoculum of 5×10^6^ CFUs was implanted into the lung via cannula. Following infection, mice were monitored twice a day for four days. Mice that lost >20% body weight and presented signs of severe clinical disease were sacrificed by CO2 administration before termination of the experiment. Futher details are outlined in supplementary materials.

### Agar-bead infection model

C57BL/6 male mice (6-10 weeks of age) were purchased from Charles-River Laboratries, Germany. The agar bead mouse model was performed according to established procedures (49). Fresh cultures were prepared in 5 ml TSB and incubated for 3 h. Bacterial cells were harvested and embedded in agar beads according to Brangonzi and colleagues (50). Five to ten mice were used for experiments and intratracheally infected with 4.6×10^6^ CFUs. Following infection, mice were monitored twice a day for 2 days. Mice that lost >20% body weight and presented signs of sever clinical disease were sacrificed by injection of 2ml 20% pentobarbital. For quantitative bacteriology, lung, liver and spleen were excised aseptically and homogenized using the homogenizer DIAX 900 (Heidolph GmbH, Schwabach, Germany). Bacterial numbers in the organs were determined by 10-fold serial dilutions of the homogenates, spotted onto blood plates after incubation at 37°C for 18 h.

### Ethics statement

Acute murine infection studies were conducted according to protocols approved by the San Raffaele Scientific Institute (Milan, Italy) Institutional Animal Care and Use Committee (IACUC) and adhered strictly to the Italian Ministry of Health guidelines for the use and care of experimental animals. Agar-bead infection studies were conducted according to protocols approved by Institute of Medical Microbiology and Hygiene (Tübingen, Germany) and adhered strictly to guidelines set by German Minstry of Health and Animal Welfare Institute (Baden-Württemberg).

### PA4130 expression and purification

The PA4130 *orf* was amplified by from PAO1 genomic DNA using primer pair NT4130F1/NT4130R1 containing N-terminal hexahistidyl tag and EcoRI/SacI restriction sites (**Table S2**). The modified PA4130 fragment was cloned into vector pSK67 using EcoRI/SacI restriction sites, resulting in plasmid pSK4130-N. The *cysG orf* was PCR amplified from *E. coli* BL21 (DE3) using primer pair CDFcysGF1/ CDFcysGR1 containing restriction sites BglII/XhoI (**Table S2**). The modified *cysG* gene was inserted into vector PCDF-DUET1 using restriction sites BglII/XhoI, producing vector pCDF-cysG. The resulting pSK4130-N and pCDF-*cysG* coding sequences were confirmed with commercial Sanger sequencing.

Vectors pSK4130-N and pCDF-*cysG* were co-transformed into *E. coli* NiCo21 by electroporation with simultaneous selection of both vectors on LB agar plates using carbenicillin and streptomycin. Single colonies selected and grown in LB broth for 16h at 37°C. Cell density was then adjusted to an OD of 0.05 into TB and further grown to an OD of ~0.6-0.8 (600nm). Cultures were then cooled to 20°C before addition of 0.05M of ferric citrate and induction with 0.1mM isopropyl β-D-thiogalactopyranoside. Following a further 18h growth cell pellets were harvested via centrifugation, flash frozen in liquid N2 and stored at −80°C.

For purification, a combination of immobilized metal ion affinity chromatography (IMAC), chitin column chromatography (CCC) used to obtain soluble PA4130. Frozen pellets were re-suspended in lysis buffer consisting of 50mM Tris-HCl, 150mM NaCl, 1.2ug/ml lysozyme and cOmplete ULTRA EDTA-free protease inhibitor cocktail at pH 8.0 with 1g cell paste per 10ml of buffer. Samples were incubated on ice for ~30 minutes with membrane disruption achieved by sonication (Fisher model 505/705).

IMAC was performed with the HiTrap chelating HP column (GE healthcare) charged with NiSO_4_. The column was equilibrated with 10 column volumes (CV) of buffer A containing 50mM Tris-HCl, 500mM NaCl, 5% glycerol, 20mM imidazole at pH 8.0 before sample application with a peristaltic pump (GE healthcare model P1). PA4130 was obtained using a linear imidazole gradient from 20-400mM imidazole between buffer A and buffer B (buffer A with 400mM imidazole) with red/brown samples collected and pooled together.

Following IMAC samples were processed and applied to chitin resin according to manufacturer’s instructions (New England Biolabs). The CCC flow-through harbouring PA4130 was collected and analysed by SDS-PAGE for homogeneity. For Size-exclusion chromatography, samples were concentrated to 5ml using a Vivaspin centrifugal filter unit (Cytiva) and injected onto a 16/600 Superdex 200pg column (Cytiva) and eluted with 50mM Tris and 150mM NaCl pH 7.5.

### Reduced methyl viologen assay

The PA4130 protein was tested for nitrite/sulphite reductase activity using the artificial electron donor methyl viologen (MV), which reduction reaction using in presence of sodium dithionite produces a blue colouration. The assay was adapted from Schnell and colleagues (25) under anaerobic conditions to prevent oxygen dependant oxidation of MV. MV oxidation was tracked spectrophotometrically at 5 second intervals with nitrite, hydroxylamine or sulphite acting as an electron acceptor.

### Griess diazotization and ammonia detection assays

Griess diazotization was performed using the Griess reagent system (Promega) according to manufacturer’s instructions. The assays were performed as above, with the exception that the electron donors were altered. MV, spinach ferredoxin and NADPH+FMN were used, with MV and ferredoxin artificially reduced using sodium dithionite. Temporal tracking of reduction reaction was achieved via sacrifice of eight independent reactions every 30 seconds.

Production of ammonia was confirmed using an ammonia assay kit (Sigma) according to manufacturer instructions.

## ACKNOWLEDGMENTS

This study was supported by European Commission (Grant NABATIVI, EU-FP7-HEALTH-2007-B, contract number 223670), Wellcome Trust AAMR DTP program (Grant 108876/Z/15/Z) and Ministero della Salute (project GR/2009/1579812). MC and PW are partly funded by the National Biofilms Innovation Centre (NBIC) which is an Innovation and Knowledge Centre funded by the Biotechnology and Biological Sciences Research Council, InnovateUK and Hartree Centre (Award Number BB/R012415/1).

We would also like to give special thanks to Dr. Annika Schmidt for her help in obtaining data from the lab of the late Professor Gerd Döring. Her efforts have been invaluable in ensuring that the work of his lab and its previous members continues to gain the recognistion it deserves.

