## Supplemental information for "NirA is an alternative nitrite reductase from *Pseudomonas aeruginosa* with potential as an anti-virulence target"

**Supplemental Materials and Methods**

**Bacteria preparation for acute infection.** An aliquot of *P. aeruginosa* strain from glycerol stocks was streaked for isolation on tryptic soy agar and incubated at 37°C O/N. Bacterial glycerol stocks were not used more than three times to avoid variability in the animal experiments. One colony was picked from the plate and used to inoculate 5 ml of tryptic soy broth (TSB) (BD, Becton and Dickinson) and placed in a shaking incubator at 37°C 200 rpm O/N. The O/N bacterial suspension was diluted to 0.1 OD/ml in 20 ml of TSB / flask and grown for 3 h at 37°C at 200rpm, to reach the log phase (1). The bacteria were pelleted by centrifugation (2,700 *g*, 15 min, 4°C), resuspended in sterile phosphate-buffered saline (PBS) and diluted to give the required dose in 60 µl (5x10^6^ CFUs).

**Mouse model of acute lung infection.** Immunocompetent C57BL/6NCrlBR male mice (8-10 weeks of age) were purchased from Charles River (Calco, Italy), shipped in protective, filtered containers, transported in climate-controlled trucks, and allowed to acclimatise for at least two days in the Animal House prior to use. Mice were maintained in the biosafety level 3 (BSL3) facility at San Raffaele Scientific Institute (Milano, Italia) where 3-5 mice per cage were housed. Mice were maintained in sterile ventilated cages. Mice were fed with standard rodent autoclaved chow (VRFI, Special Diets Services, UK) and autoclaved tap water. Fluorescent lights were cycled 12h on, 12h off, and ambient temperature (23±1°C) and relative humidity (40-60%) were regulated.

For infection experiments, mice were anaesthetized by an intraperitoneal injection of a solution of Avertin (2,2,2- tribromethanol, 97%) in 0.9% NaCl and administered at a volume of 0.015 ml/g body weight. Mice were placed in supine position. The trachea was directly visualised by ventral midline, exposed and intubated with a sterile, flexible 22-g cannula attached to a 1 ml syringe. An inoculum of 60 μl of planktonic bacterial cells was implanted via the cannula into the lung. After inoculation, all incisions were closed by suture. Infections were all performed in the late morning. In addition, in all the experiments, mice had been subdivided according to the bodyweight to have similar mean in the two groups of infection (PAO1-L versus PAJD25).

Mice were monitored twice a day for coat quality, posture, attitude, ambulation, hydration status and body weight. Mice that lost >20% body weight and had evidence of severe clinical disease, such as scruffy coat, inactivity, loss of appetite, poor locomotion, or painful posture, were sacrificed before the termination of the experiments with an overdose of carbon dioxide. At day 4 post-infection, surviving mice were sacrificed with an overdose of carbon dioxide.

Animal studies were conducted according to protocols approved by San Raffaele Scientific Institute (Milan, Italy) Institutional Animal Care and Use Committee (IACUC) and adhered strictly to the Italian Ministry of Health guidelines for the use and care of experimental animals.

**Supplemental Data**

**Table S1** Twitching, swimming, protease and elastase production in PA4130 interrupted strains.

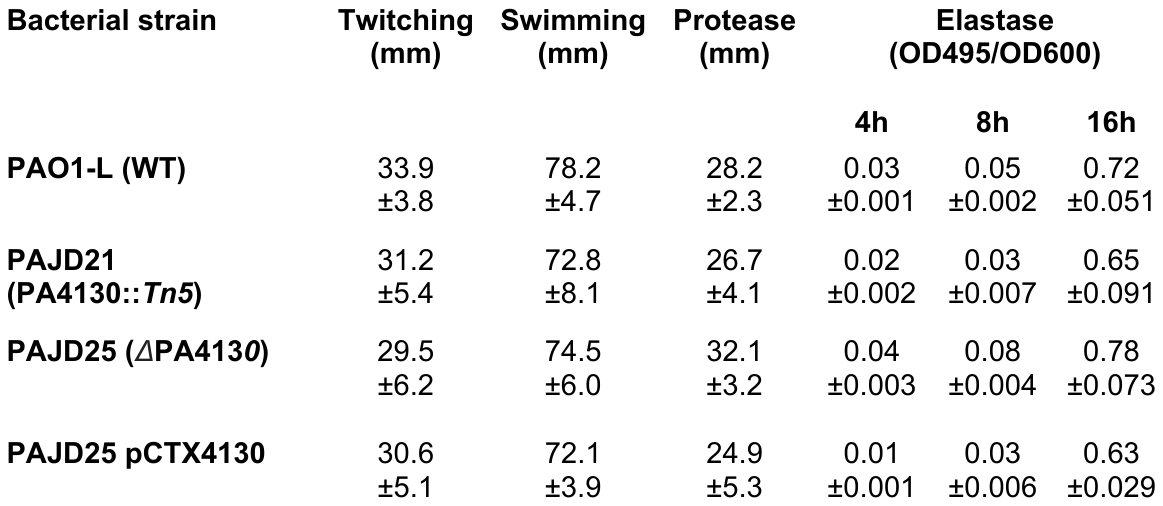

**Table S2** Bacterial strains, plasmids and primers used in this study.

| **Strains** | **Description** | **Origin** |
| --- | --- | --- |
| ***P. aeruginosa* strains** |  |  |
| PAO1-L | PAO1 Lausanne collection wild type | (2, 3) |
| PAJD21 | Strain PAO1-L with Tn5 insertion in PA4130 Gm^R^ | This study |
| PASF06 | In-frame marker-less deletion of PA4129 | This study |
| PAJD25 | In frame marker-less deletion of PA4130 | This study |
| PA7 Bo599 | Clinical PA7 strain | (4) |
| PA7 Bo599 ΔPA4130 | In-frame marker-less deletion of PA4130 orthologue | This study |
| PA14 AUS471 | Clinical PA14 strain | (4) |
| PA14 AUS471 ΔPA4130 | In-frame marker-less deletion of PA4130 orthologue | This study |
| LESB58 PA-W39 | Clinical LESB58 strain isolated from wound. | (4) |
| LESB58 PA-W39 ΔPA4130 | In-frame marker-less deletion of PA4130 orthologue | This study |
| ***E. coli* strains** |  |  |
| NEB5-alpha | F−,φ80dl*acZ*ΔM15,Δ(*lacZYA-argF*)U169,*deoR*,*recA1*, *endA1*, *hsdR17*(rk−,mk+), *phoA*, *supE44*, λ−, thi1, *gyrA96*, *relA1* | New England Biolabs |
| S17.1 λpir | pro, res^−^ *hsdR17* (rK^−^ mK^+^) *recA^−^* with an integrated RP4-2-Tc::Mu-Km::Tn7, Tp^r^ *λpir* | (5) |
| BL21 (DE3) | F– *ompT* *gal* *dcm lon* *hsdSB*(rB–mB–) λ(DE3 [*lacI* lacUV5-T7p07 *ind1* *sam7* *nin5*]) [*malB+*]K-12(λS) | (6) |
| NiCo21 (DE3) | F– *ompT* *gal dcm lon hsdSB*(rB–mB–) λ(DE3 [*lacI* lacUV5-T7p07 *ind1* *sam7* *nin5*]) [*malB+*]K-12(λS) *glmS6ala* *slyD-CBD* *arnA-CBD* | (7) |
| **Plasmids** |  |  |
| pME3087 | Suicide vector, ColE1 replicon, Tc^R^ | (8) |
| pME4129 | pME3087 based vector with upstream and downstream regions of PA4129 spliced together for markerless deletion generation. | This study |
| pME4130 | pME3087 based vector with upstream and downstream regions of PA4130 spliced together for double-crossover generation | This study |
| Mini-CTX-1 | *aatB* P. aeruginosa integrative vector, Tc^R^ | (9) |
| pCTX4130 | Integrative PA4130 complementation vector under control of the native promoter (+498bp), Tc^R^ | This study |
| pSK67 | pTOPO type vector for protein overexpression. Under control of a T7 promoter and IPTG inducible. Amp^R^ |  |
| pSK4130 | pSK67 based vector with N-terminal hexahistidyl tagged PA4130 inserted at the EcoR1 and SacI restriction sites for overexpression | This study |
| pCDF-DUET1 | DUET vector containing 2 T7 promoters under the control of LacI. Enables co-expression of up to 4 target proteins. Sp^R^ | Novagen |
| pCDF-*cysG* | pCDF-DUET1 based vectors with *cysG* inserted into MCS2 at the NcoI and XhoI site for overexpresssion | This study |
| **Primers** | **Sequence** | **Modifications** |
| 4129DELF1 | 5’-ATAGAATTCTGTGGCGCGAGGCCTGCG-3’ | EcoRI |
| 4129DELR1 | 5’-CTAGCGTCGGCGGAACAGGTTGTTCATGCCGGTTCC-3’ | N/A |
| 4129DELF2 | 5’-GGCATGAACAACCTGTTCCGCCGACGCTAGGCATAC-3’ | N/A |
| 4129DELR2 | 5’-TATGGATCCGCTGGAACAGCGTGGCGGAG-3’ | BamHI |
| 4130DELF1 | 5’-ATATCTAGATCATTTTTCGTAGGCCCATC-3’ | XbaI |
| 4130DELR1 | 5’-TCATGCCGGTTCCTCGTACTGGTACATCGCAAAGCC-3’ | N/A |
| 4130DELF2 | 5’-GCGATGTACCAGTACGAGGAACCGGCATGAACAACC-3’ | N/A |
| 4130DELR2 | 5’-ATAAAGCTTTCCTCGACGTTCTTGTCCTC-3’ | HindIII |
| 4130CTXF1 | 5’-ATAAAGCTTGGGCCGTTCACCGCCGAC-3’ | HindIII |
| 4130CTXR1 | 5’-TATGGATCCTCATGCCGGTTCCTCCATCCTG-3’ | BamHI |
| NTH-30F1 | 5’TTAGAATTCATGCATCACCATCACCATCACTACCAGTACGATGAATACG-3’ | EcoRI + 6XHis |
| NTH-30R1 | 5’-TTACTCGAGTCATGCCGGTTCCTCCATCCT-3’ | SacI |
| M2cysGF1 | 5’-ATACCATGGGTGGATCATTTGCCTATATTTTGC-3’ | NcoI |
| M2cysGR1 | 5’-TATGGATCCTTAATGGTTGGAGAACCAGTTCAG-3’ | XhoI |

| **Assay** | **PASF06** | **PAJD25** |
| --- | --- | --- |
| Pyocyanin | 88±10 | 48±7** |
| Pyoverdine | 94±12 | 67±10* |
| Swarming surface coverage | 106±7 | 20±4**** |
| Elastase | 91±7 | 108±5 |
| Protease | 90±9 | 92±5 |

**Table S3** Virulence factor production from strains PASF06 (ΔPA4129) and PAJD25 (ΔPA4130) expressed as a percentage (%) of the wild type strain PAO1-L. Data collected from at least 2 separate experiments with 3 to 5 replicates.

**Table S4** Virulence factor production from PA4130 orthologue mutant strains expressed as a percentage (%) of the wild-type strain at 20H. Data collated from 2 separate experiments with 3 to 5 replicates. Red text denotes enhanced phenotype.

| **Assay** | **PA7 Bo599 PA4130** | **PA14 AUS471 PA4130** | **LESB58 PA-W39 PA4130** |
| --- | --- | --- | --- |
| Pyocyanin | 54.7±10**** | 52.7±3.5**** | 44.9±9.3**** |
| Pyoverdine | 140±6**** | 53.7±2.5 | 206±31**** |
| Swarming surface coverage | 56.2±9.2**** | 12.7±2.5**** | 42.3±8.5* |
| Protease | 110±12.9 | 89.7±9.1 | 111.1±6.6 |

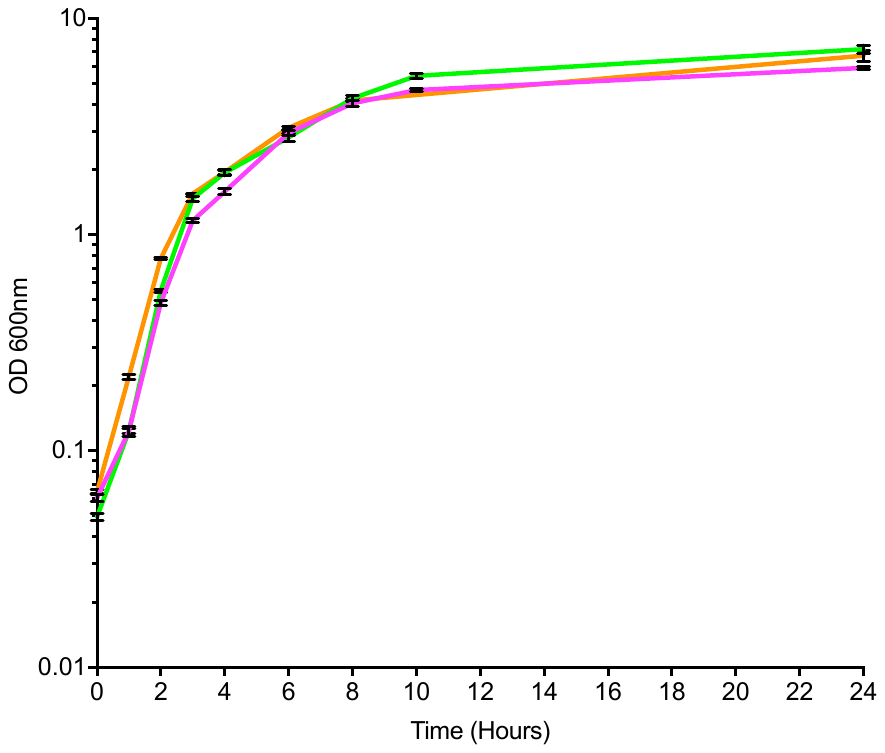

**A**

Optical density (600nm)

PAO1-L

PAJD25

PASF06

Time (Hours)

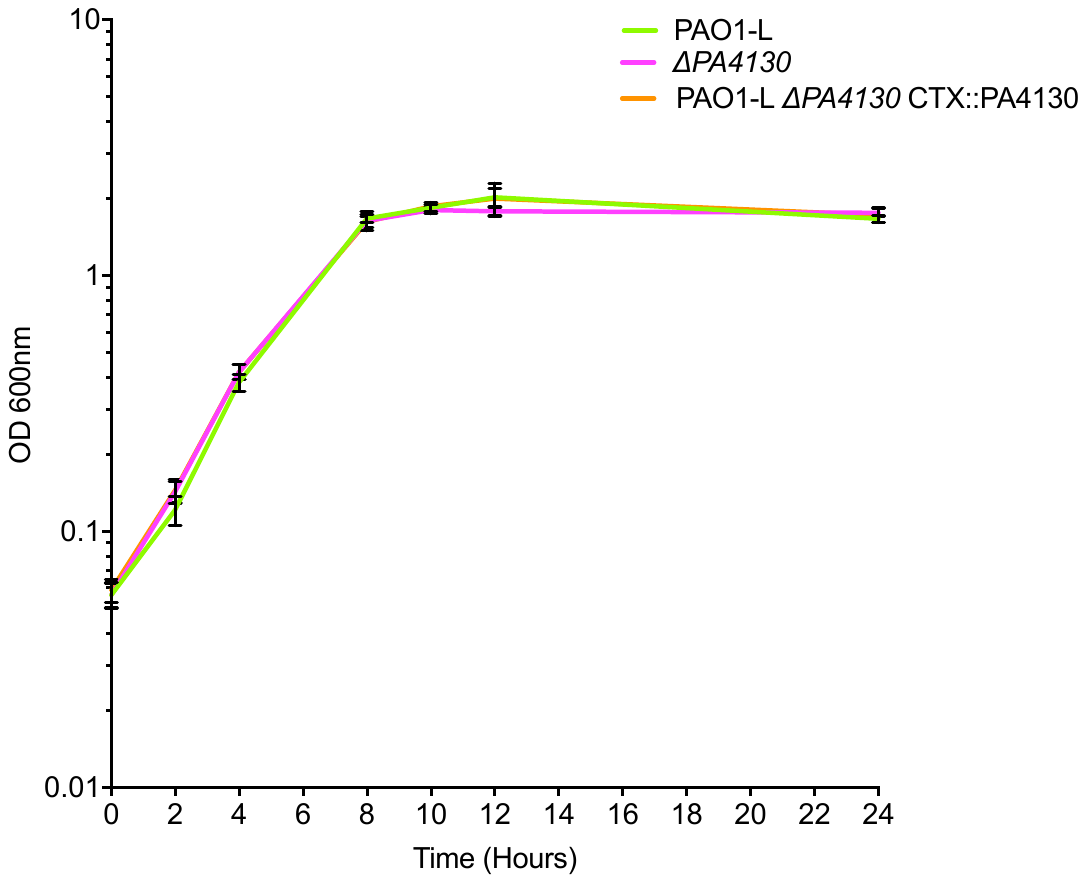

**BA**

Optical density (600nm)

PASF06

PAJD25

PAO1-L

Time (Hours)

**FIG S1** Growth curves of PAO1-L, PAJD25 (ΔPA4130) and PASF06 (ΔPA4129) in (A) LB and (B) Modified artificial sputum media demonstrating no growth defect of the mutants when compared to WT.

```
EcoliCysI      ---------------------------------------------------------MSE
SpinachNirA    MASLPVNKIIPSSTTLLSSSNNNRRRNNSSIRCQKAVSPAAETAAVSPSVDAARLEPRVE
PA4130         -------------------------------------------------------MYQYD
M_tubSirA      ---------------------------------------------------MTTARPAKA

EcoliCysI      KHPGPLVV----EGKLTDAERMKHESNYLRGTIAEDLNDGLTGGFKGDNFLLIRFH----
SpinachNirA    ERDGFWVLKEEFRSGINPAEKVKIEKDPMKLFIEDGISDLAT--LSMEEVDKSK-HNKDD
PA4130         EYDQALV-----------SERVAQFRD--------QIARRLDGELSEEEFLPLRLQN---
M_tubSirA      RNEGQWAL--GHREPLNANEELKKAGNPL------DVRERIENIYAKQGFDSI---DKTD

EcoliCysI      --------GMYQQDDRDIR----AERAEQKLEPRHAMLLRCRLPGGVITTKQWQAIDKFA
SpinachNirA    IDVRLKWLGLFHRRKHHYG----R------------FMMRLKLPNGVTTSEQTRYLASVI
PA4130         --------GLYLQKHA--------------------YMLRVAIPYGTLSAPQLRALAHVA
M_tubSirA      LRGRFRWWGLYTQREQGYDGTWTGDDNIDKLEAKY-FMMRVRCDGGALSAAALRTLGQIS

EcoliCysI      GENTIYGSIRLTNRQTFQFHGILKKNVKPVHQMLHSVGLDALATANDMNRNVLCTSNPYE
SpinachNirA    KKYGKDGCADVTTRQNWQIRGVVLPDVPEIIKGLESVGLTSLQSGMDNVRNPV--GNPLA
PA4130         RHYDR-GYGHFTTRQNIQFNWIELEQVGDILEHLAGAQMHAIQTSGNCVRNIT--TEAFA
M_tubSirA      TEFAR-DTADISDRQNVQYHWIEVENVPEIWRRLDDVGLQTTEACGDCPRVVL--GSPLA

EcoliCysI      SQLHAEAYEWAKKISEHLLPRTR-AYAEIWLDQEKVATTDEEPILGQTYLPRKFKTTVVI
SpinachNirA    GI---DPHEIVD---------TR-PFTNLI---SQFVTANSRGNLSITNLPRKWNPCVIG
PA4130         GV---AADEWTD---------PR-PLAEIL---RQWSTVNP----EFLFLPRKFKIALSS
M_tubSirA      GE---SLDEVLD---------PTWAIEEIV---RRYIG-KP----DFADLPRKYKTAISG

EcoliCysI      PPQNDIDLHANDMNFVAI--AENGKLVGFNLLVGGGLSIEHGNKKTYARTASEFGYLPLE
SpinachNirA    SHDLYEHPHINDLAYMPA--TKNGKF-GFNLLVGGFFSIKRCEEAIPL-----DAWVSAE
PA4130         AVEDRAAVQMHDIGLYLYRHPDAGEL-RLRVLVGGGLG------RTPMLGQVIRDDLPWQ
M_tubSirA      LQD--VAHEINDVAFIGVNHPEHGP--GLDLWVGGGLS------TNPMLAQRVGAWVPLG

EcoliCysI      HTLAVAEAVVTTQRDWGNRTDRKNAKTKYTLERVGVETFKAEVERRAGIKFEPIRPY---
SpinachNirA    DVVPVCKAMLEAFRDLGFRGNRQKCRMMWLIDELGMEAFRGEVEKRMPEQVLERASS---
PA4130         HLLSYVEAILRVYNRYGRRDNKYKARIKILVKALGIEAFAREVEEEW--QHLRDGPAQLT
M_tubSirA      EVPEVWAAVTSVFRDYGYRRLRAKARLKFLIKDWGIAKFREVLETEYLKRPLIDGPA---

EcolicysI      ----------------------------------EFTG----RGDRIGWVKGIDDNWHLT
SpinachNirA    ---------------------------------EELVQKDWERREYLGVHPQKQQGLSFV
PA4130         AEECQRVAERFVLPRYLPPADGELAYGSARAADPAFAR--WASR-NVQAHK-VPGYASVV
M_tubSirA      ---------------------------------PEPVK---HPIDHVGVQR-LKNGLNAV

EcolicysI      LFIENGRILDYPARPLKTGLLE-----IAKIHKGDFRITANQNLIIAGVPESEKAKIEKI
SpinachNirA    -----G--LHIPVGRLQADEMEELARIADVYGSGELRLTVEQNIII---PNVENSKIDSL
PA4130         LSTKPG--ASAPPGDVTAEQMERVADWAERYGFGEIRVAHEQNLVL---PDVRLENLHAL
M_tubSirA      -----G--VAPIAGRVSGTILTAVADLMARAGSDRIRFTPYQKLVILDIPDALLDDLIAG

EcolicysI      AKESGLMNAVTP----QRENSMACVSFPTCPLA----MAEAERFLPSFIDNIDNLMAKHG
SpinachNirA    LNEPLLKERYSPEPPILMKGLVACTGSQFCGQAIIETKARALKVTEEVQ-RLVSVTR---
PA4130         WREACAAGLGTPNQG-LLSDIIACPGGDYCALA----NAKSIPIAQGIQQRFEDLDHLHD
M_tubSirA      LDALGLQSRPSH----WRRNLMACSGIEFCKLSFAETRVRAQHLVPELERRLEDINSQLD

EcolicysI      VSDEHIVMRVTGCPNGCGRAMLAEVGLVG----KAPGR----YNLHLGGNR-IGTRIPRM
SpinachNirA    --P--VRMHWTGCPNSCGQVQVADIGFMGCMTRDENGKPCEGADVFVGGRIGSDSHLGDI
PA4130         IGE--LSLNISGCMNACGHHHIGNIGILGV---DKSGS--EWYQVTLGGAQGKDSALGKV
M_tubSirA      V-P--ITVNINGCPNSCARIQIADIGFKGQMIDDGHGGSVEGFQVHLGGHLGLDAGFGRK

EcolicysI      YKE-NITEPEILASLDELIGRW---AKEREAGEGFGDFTVRAGI--IRPVLDPARDLWD
SpinachNirA    YKKAVPCKDLVPVVAEILINQFGAVPREREEAE--------------------------
PA4130         IGP-SFSAAEVPAVIERIVETF---TDLRVGPERFIDTFNRVGLEPFKARVYARMEEPA
M_tubSirA      LRQHKVTSDELGDYIDRVVRNF---VKHRSEGERFAQWVIRAEEDDLR-----------
```

**FIG S2** Amino Acid alignment performed with ClustalW comparing PA4130 with homologous proteins CysI (*E. coli*), NirA (Spinach) and SirA (*M. tuberculosis*). Highlighted in green are Siroheme interacting residues observed in the crystal structures of CysI, NirA and SirA with conserved residues of PA4130 highlighted in pink. Conservation of these residues along with four essential cysteine residues required for iron-sulfur cluster insertion suggests that PA4130 requires Siroheme and 4Fe-4S for its functional activity.

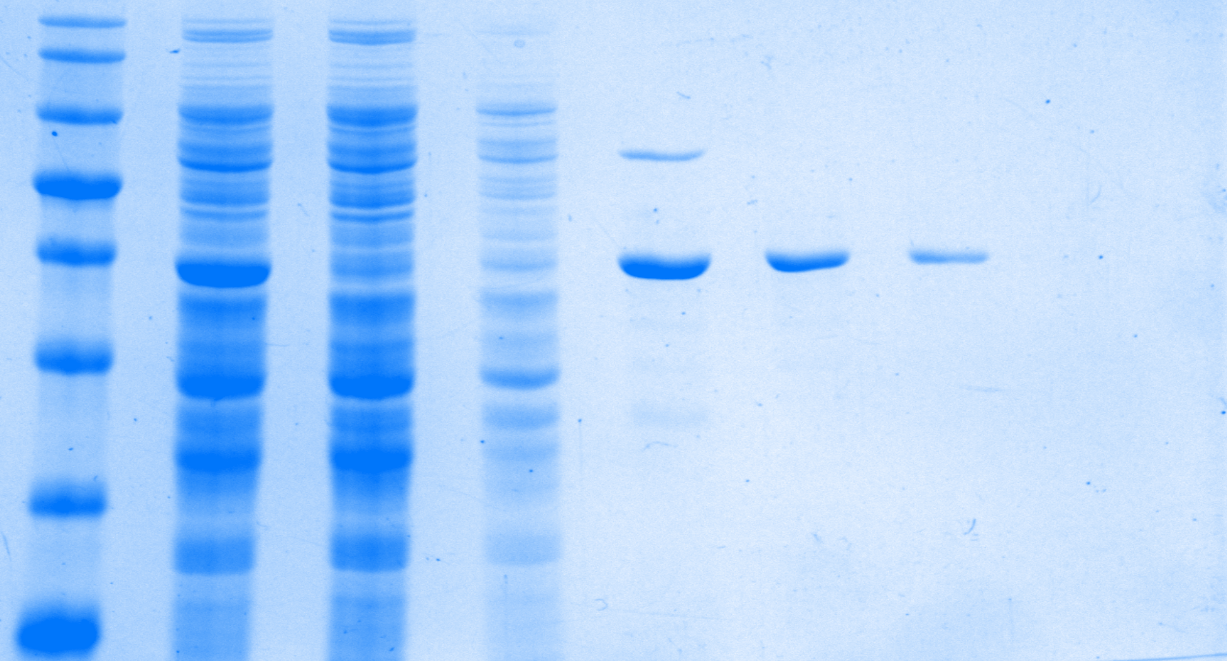

**kDa	1	2	3	4	5	6	7	8	9	10	11	12	13**

**58**

**46**

**38**

**25**

**80**

**100**

**135**

**PA4130**

**CysG**

**B**

**A**

Absorbance (280nm)

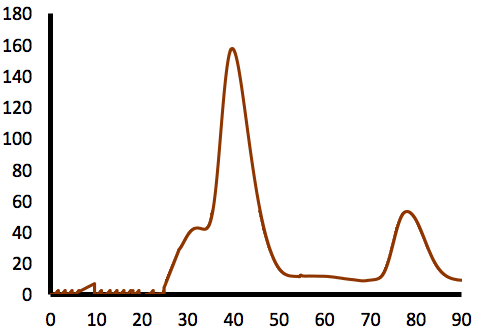

Volume (ml)

**Fig S3** PA4130 purification using *E. coli* NiCo21 pSK4130-N/pCDF-*cysG*.(A) Protein purification was performed with a combination of immobilised metal ion chromatography (IMAC) and chitin column chromatography (CCC). Lane 1- broad-range MWt ladder; lane 2- Soluble lysate fraction; lane 3- lysate IMAC flow-through; lane 4- IMAC 40mM imidazole wash; lane 5- post IMAC PA4130 sample; lane 6- CCC flow-through; lane 7-CCC wash.(B) Size exclusion chromatogram displaying elution profile of PA4130 post-NINTA. PA4130 flows as a monomer with a predicted molecular weight of 61kDa.

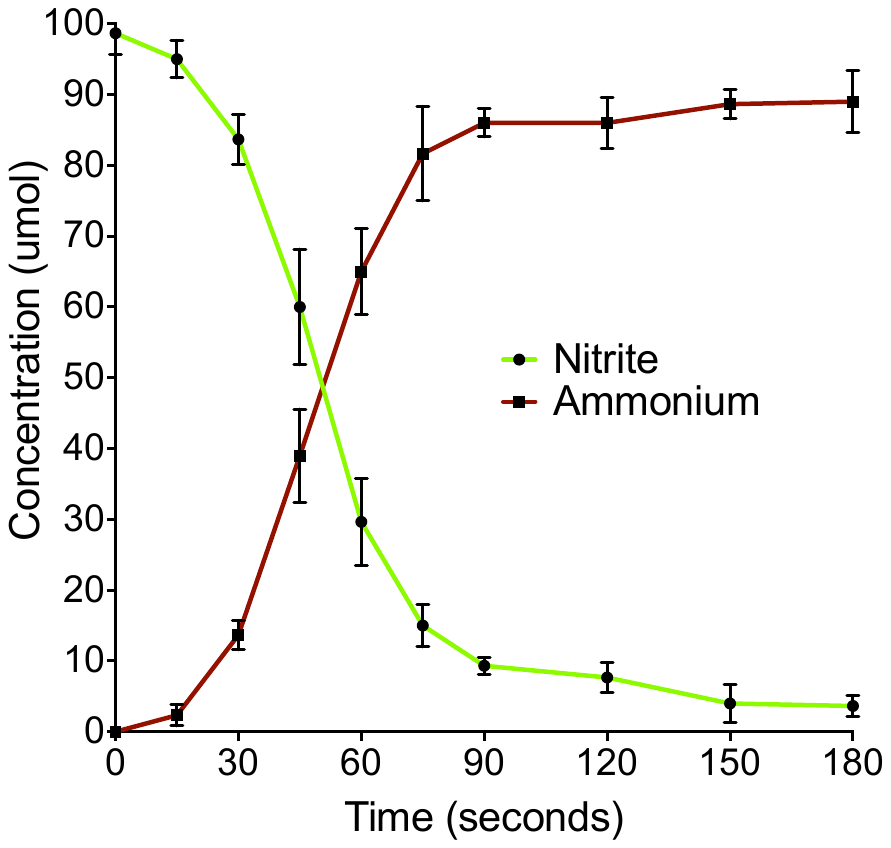

Concentration (micro-mol)

Time (seconds)

**Fig S4** Detection of remaining nitrite and ammonium production in a reduced ferredoxin dependant nitrite reduction assay with PA4130. Nitrite levels decrease concomitantly with an increase in ammonium production at a 1:1 ratio. Nitrite and ammonium concentrations were determined using Griess diazotisation and an ammonia assay kit. Concentrations were calculated through interpolation of absorbance values on standard curves prepared with KNO_2_ and NH_4_Cl over a range of 0-120umol.
